# Moderated designs can balance between batch-effect mitigation and cell loss due to hashtag-assisted pooling in single-cell experiments

**DOI:** 10.1101/2025.10.16.682943

**Authors:** Budha Chatterjee, Katrina Gorga, Carly Blair, Yuko Ohta, Elizabeth M. Hill, Christopher T. Boughter, Martin Meier-Schellersheim, Nevil J. Singh

## Abstract

Minimizing experimental noise is integral to robust data generation in single-cell science. Experimental processing of different samples as a single pool, made possible by hashtag-assisted pooling, helps minimize batch-effects, but the computational demultiplexing of the data can also lead to loss of cells whose hashtags cannot be resolved accurately. Here, we examine four alternate experimental designs that could be used instead of a single-pool approach and quantify the batch effects as well as cell loss in each case. While a reference design offers the best performance, this study can help individual investigators choose one suited for their biological questions.

## Background

Single-cell RNAseq (scRNA-seq) methodologies involve significant amplification of transcripts in each cell during the preparation of libraries, which can amplify well-to-well variability stemming from technical rather than biological reasons - commonly referred to as ‘batch-effects’ [1-4]. Sample multiplexing, whereby cells from different samples in the same experiment are individually barcoded (i.e., hash-tagged) and then combined in the same well, circumvents this problem [5-11] (Fig. 1A). The benefit of sample multiplexing is not just in the mitigation of batch-effects, but also a reduction in per-sample cost since this allows an investigator to analyze fewer cells per sample in a single well, rather than dedicating the maximum capacity of each well to just one sample.

**Figure 1.**
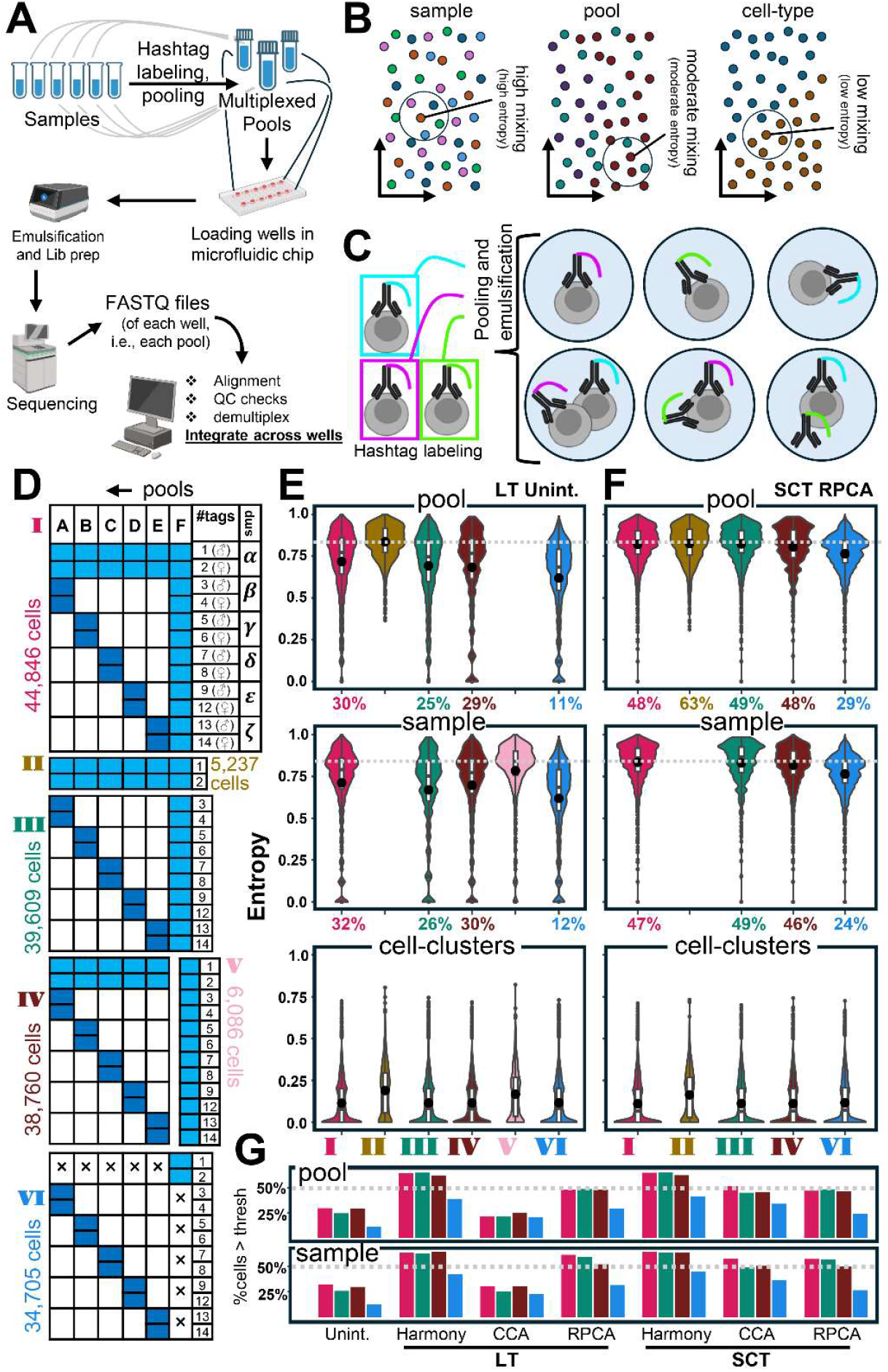
A cartoon overview of single-cell measurements and estimation of batch effects in different designs of the wells pre- and post-data integration. (A) Overview of an in-house, single-day, scRNA-seq experiment in a laboratory. Samples are labeled with hashtags and combined for multiplexing. Each multiplexed pool is loaded to separate wells on a microfluidic plate (e.g. 10xGenomics chips) for droplet capture and reaction. The libraries are sequenced, and the reads from each well (i.e., pool) separated based on the indexes they receive during library preparation. FASTQ files are then taken through the computational pipelines of alignment, quality control, demultiplexing etc. Finally, the data from all the wells in that experiment are integrated to mitigate batch effects. (B) In scRNA-seq data analysis pipelines, the cells are projected in lower-dimensional spaces (e.g., principal component space or further corrected integrated spaces) based on their transcriptome. Generally, 30-50 dimensions are used but, in the cartoon shown here, a two-dimensional space is depicted for illustration. Each cell can have different attributes, also known as metadata, as depicted by different colors here: sample (left), pool (middle), cell-type (right). In the absence of batch-effects, similar cells from different pools and samples will be well-mixed in this space, since they will not have batch-related differences in their gene expression, other than biologically relevant differences. The lower the batch effects, the higher the diversity of the neighborhood of a cell and the larger is the entropy of that cell for that metadata. Typically, cell-type metadata has low mixing compared to that of samples and pools, since cells of the same type are expected to remain together in the reduction space. (C) A cartoon of emulsion droplets with hashtag labelled cells. Generally, they are individual cells with one predominant hashtag (top circles). In some cases, more than one cell can be in a droplet (left bottom circle). Or one cell may be labelled by more than one hashtag due to ambient hashtag antibodies after pooling (middle bottom). Or simply the process of emulsification might capture free antibodies in the medium (right bottom). (D) Different designs of data are considered for batch-effect calculations. Columns in each block indicate wells (A to F), rows indicate samples (α to ζ, and hashtags: 1 to 9, 12 to 14 [10 and 11 is skipped due to QC considerations, see Methods]). Each design is indicated by colored boxes in each block. The lighter blue boxes indicate four times higher loading of cells than those in navy blue. Number of cells in each design and the design names in Roman numerals are shown in colors matching those used in parts (E) and (F). Design-I (top) illustrates the compound format used in the entire experiment. Descriptions of the designs II to VI are given in the text. (E) Entropies of each cell in different designs are measured, and the distributions of the entropies are presented as violin plots. Entropies of pools (top), samples (middle) and cell-clusters (i.e., cell-types) (bottom) are shown. For design-V and design-II, pool and sample entropies, respectively, are not shown (see text, Methods). The percentages in the X-axis are %cells above the threshold (dashed gray line) that was chosen as the median of the pool-entropy for design-II (top) or that of sample-entropy in design-V (middle). Unint: unintegrated; LT: log transformed. (F) Same as (E) but looking at the mixing in SC-transformed RPCA (see text) integrated reduced space. Design-V is excluded in this representation since this design has only one well, hence cannot be integrated. SCT: SC-transformed. (G) Percentage of cells above the pool and sample entropy thresholds obtained with three different transformation/integration strategies (see text, Methods) for designs I, III, IV and VI.

One of the widely used metrics to measure batch-effects, borrowed from information theory, is entropy [2, 4] (normalized Shannon entropy, Fig. 1B, Methods). When individual cells are projected in lower-dimensional space based on their transcriptome, biologically similar cells from different ‘batches’ should be randomly mixed in the neighborhood of a given cell in the latter space. For each cell, its entropy is the measurement of the numeric mixing of its neighboring cells from different ‘batches’.

Hashtag multiplexing itself has the potential to introduce noise (Fig. 1C). Apart from encapsulation of multiple hash-tagged cells or ambiently available hashtag antibodies during microfluidic emulsification, there can be exchange of hashtags between cells after pooling, since the hashtags are against the same antigens [11, 12]. All reads from some of these latter droplets (bottom circles, Fig. 1C) will be discarded during computational demultiplexing. Thus, all hashtag-multiplexing processes are inherently lossy to different extents [7, 13]. To our knowledge, a systematic evaluation of the batch-effects and cell loss in different experimental design strategies that use hashtag-multiplexing, has not yet been performed.

Here, we perform a single compound experiment using the 10xGenomics scRNA-seq pipeline to then computationally extract four major single-cell experimental designs, i.e., single-pool, reference, chain, and confounded, to quantify batch-effects (using entropy) as well as cell loss due to hashtag demultiplexing. Our analysis offers a guide towards choosing the one suitable for different experimental contexts.

## Results and Discussion

To avoid variations in sample handling across different experimental designs, we performed a single wet-lab experiment with a compound design and then extracted different designs from it for comparisons. Since the biology of the control and test groups themselves is not relevant for our analysis, we refer to these as pools of six different samples: α-ζ in different combinations in six different wells: A-F (Methods, Fig. 1D-I). Sample-α is the baseline or control group, and the rest of the samples are experimental groups. All pools received sample-α. Pools A-E received only one more sample other than sample-α. Pool F received all samples. Each sample was a 50:50 mix of female and male mice, and each received unique hashtags before pooling.

After demultiplexing (Fig. 1A, Methods) we extracted six designs from the whole experiment, which we denoted as a ‘compound’ design (Fig. 1D, design-I). Design-II included all α samples from all wells (Fig. 1D-II). Next, we removed the α samples from the design-I – to form a ‘reference’ design (Fig. 1D-III) as one pool contains all samples to serve as a ‘reference’ and rest of the pools have only some of the samples [14]. Removing pool-F from design-I yielded ‘chain’ design – where pools share certain samples (sample-α in this case), without a ‘reference’ pool (Fig. 1D-IV). Design-V has only pool-F with all samples (Fig. 1D-V). Lastly, design-VI, lacks sample-α from wells A-E and samples β-ζ from well-F and is the ‘confounded’ design, keeping one sample per well. Next, we measured batch-effects in each design after transformation (log-transform and SC-transform, Methods) and integration (Harmony, CCA, RPCA, Methods)

The design with the highest median pool-entropy, i.e., least batch-effect (for pool), before integration was design-II (median entropy= 0.842, that of design-I= 0.772, design-IV= 0.767, Fig. 1E, top), consistent with previous analysis of nuclear hashing experiments [10]. We further validated this by evaluating the differentially expressed transcripts between sample-α from pool-A vs rest of the pools B-F (Fig. S3A). We observed very minor (1 disparate upregulated DEG in three comparisons, 11 and 13 DEGs in the rest), mostly nonoverlapping (3 common genes among the latter) gene expression differences across five DEG comparisons and none were significantly enriched for any biological function (Fisher’s test, cutoff p-value = 0.01). As a positive control, we compared the samples α vs β, in pool-A and in pool-F. For these between-sample comparisons, in contrast with the within-sample comparisons (Fig. S3A), 55 and 97 genes were upregulated, respectively, among which 47 were common (Fig. S3B). We used the median of the pool-entropy for design-II as a threshold (dashed line, Fig. 1E, top panel) and checked the percentage of cells in other designs that are above it. Although approximately 25-30% of cells in designs I, III, and IV were above the threshold, the confounded design elicited a mere 11% of cells above the latter.

In the sample-entropy calculations, the unintegrated design-V showed the highest performance (median entropy= 0.839, that of design-I= 0.791, design-IV= 0.771, Fig. 1E, middle panel), as expected, since this is a reference pool where all the samples are processed together in the same well. We therefore used the median of the sample-entropy distribution of design-V as the sample-entropy threshold (dashed line, Fig. 1E, middle panel) to check the sample ‘batch’ effects in the other designs. The rationale here is that any pipeline of transformation/design/integration that would yield 50% or more cells above the latter entropy threshold is performing as well as the ‘biological’ sample mixing (i.e., pool-F – design-V). Using this criterion, we find that in designs I, III and IV, 26-32% of the cells were above the sample-entropy threshold. Again, design-VI was the worst performer, with only 12% cells above the threshold. The cell-cluster entropy distributions showed much lower medians (range of medians of entropy for designs I-VI: 0.077-0.191, Fig. 1E, bottom panel), as we would expect.

Calculating the entropy distributions post-integration (Fig. 1F, S4) using a representative transformation/integration approach (SC-transform, RPCA integration) we found that designs I, III, and IV consistently reached ∼50% of cells above the respective thresholds of pool- and sample-entropy distributions. However, the integration process could not recover the batch-effects of design-VI for either sample or pool (Fig. 1G). All integration methods in both transformations brought to the desired 50% cells above the entropy threshold, except for the log-transformed CCA. To conclude, no integration method could fully recover the batch effects in design-VI – i.e., designs III and IV always performed better than design-VI, irrespective of the subsequent computational pipeline, emphasizing that some designs are always better than others.

We then evaluated the labeling efficacies of the hashtags used. The droplet-doublets (Fig. 1C, bottom-left circle) often tend to have higher counts and features than the singlets [15, 16]. We observed that the distributions of the doublets are largely overlapping with those of singlets, i.e., ∼50% counts or features in doublets are below the 3^rd^-quartile threshold in singlets (Fig. S5A-B). This suggests that a large percentage of hashtag-doublets (Fig. 1C, bottom-, middle, and right circles) are indeed droplet-singlets [11, 12].

To measure the global efficacy of hashtags, we calculated the percentage of total number of singlets against the total number of cells, within a pool (Fig. 2A, global demultiplexing efficacy, GDE). The closer this metric is to 100%, the more successful the entire pooling and demultiplexing would be. For scenarios using 2 and 4 hashtags (pools A-E), the GDE was 93% and 90% respectively. But for pool-F, it falls abruptly to 76%, indicating that with a higher number of hashtags, more cells are likely lost as non-singlets. Next, we calculated individual hashtag efficacy (IHE), which is the percentage of singlets for a given hashtag against the total number of cells that are positive for that hashtag (Fig. 2B). IHE for the hashtags in pools A-E showed a median of ∼80%. Again, in pool-F, there was a sharp fall of the IHE. Clearly, the efficacy of the hashtags was dependent on how many of them were being used simultaneously.

**Figure 2.**
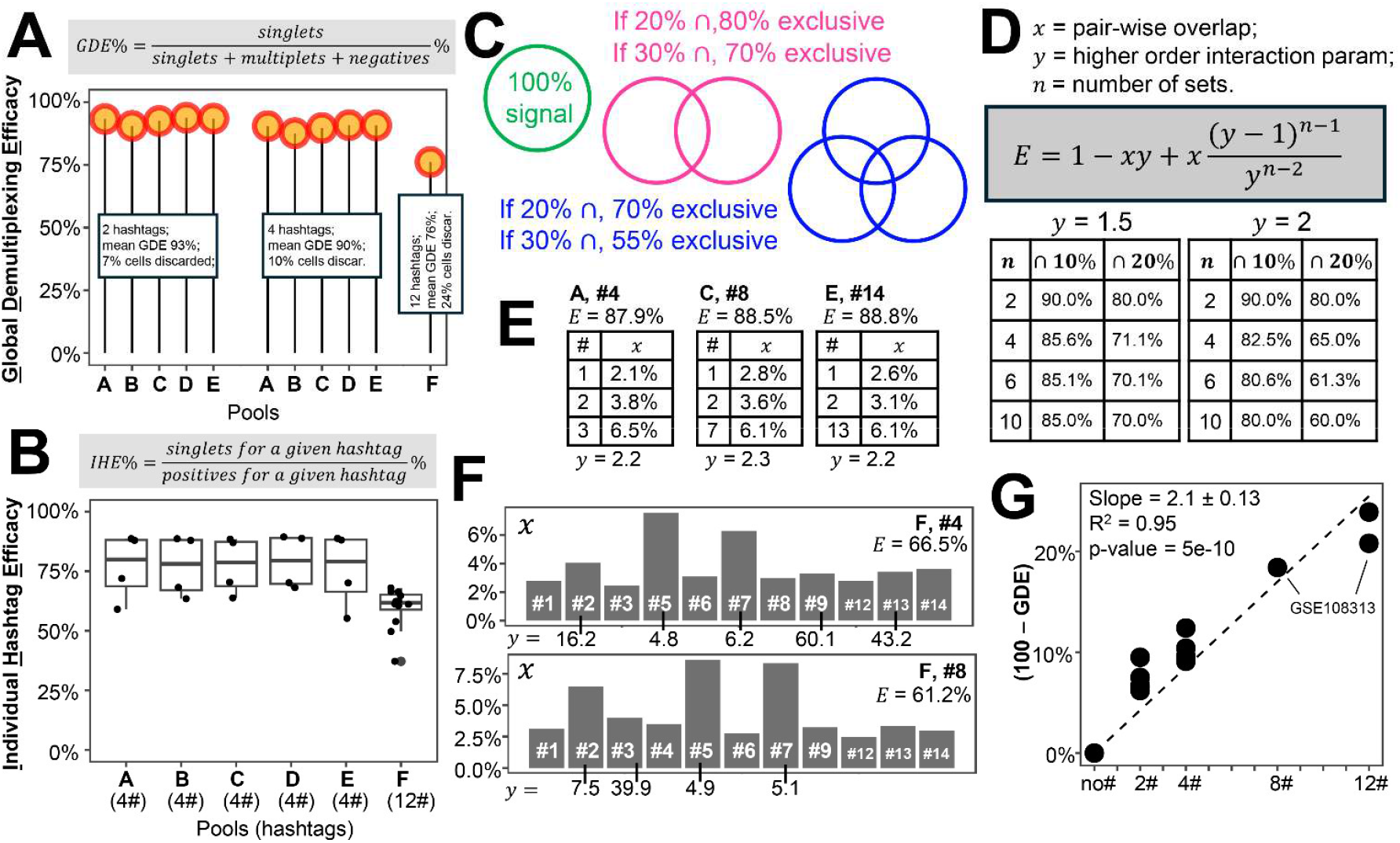
Evaluation of the performance of hashtags in demultiplexing: (A) Evaluation of global demultiplexing efficacy (GDE) in each pool. The percentage of hashtag-singlets within all cells in a well is depicted with a lollipop plot. First five lollipops are the calculations of GDEs of wells A to E, combining two hashtags within a sample as one. The approximated mean GDE is noted on the box covering the stems of the lollipops. (B) Evaluation of individual hashtag efficacies (IHE) for each hashtag used in each pool is shown as dot plots accompanied by overlaid box plots. Pools and the number of hashtags (in brackets) used in them are marked on the X-axis. The hash (#) signs are an abbreviation for ‘hashtag’. For (A) and (B), the expressions of GDE and IHE, respectively, are noted in the gray boxes at the top. (C) Cartoon Venn diagrams to illustrate the relations between overlaps and the availability of exclusive parts of intersecting sets. (D) A closed-form expression to calculate exclusive parts of sets (*E*), given that we know *x, y*, and *n* (see text, Text S1) in the dark gray box on the right. The two charts below the gray box are calculations of *E*, for the given value of *y* (shown at the top of the charts). The different values of *x* are at the top row, followed by ∩ (intersection) sign and *n* is the first column in the charts. (E, F) Estimation of *x* and *y* in single-cell data. Three charts are for the pools with four hashtags – the first columns indicate the specific hashtag for which the overlap is calculated, and the second columns are the estimated values of *x*. (E). Two bar plots at the bottom are for pool F, i.e., 12 hashtags (F). The title above the charts/bar plots, such as ‘F, #4’, indicates that *x* and *y* are estimated for hashtag 4 in well F. Below that, the percentage of exclusive signals, i.e., singlets for that hashtag, is noted. The bars or the second columns of the charts are the values of *x*, as estimated directly from data (see Methods). Numeric estimates of *y* are noted immediately below the bars or the chart entries (i.e., the corresponding *x* values). (G) The difference between GDE and 100 (as percentages) is plotted in the Y-axis for different pools, while the number of hashtags in respective pools in the X-axis, from the current dataset and another external dataset (GSE108313) [7]. Slope, p-value, and R^2^ of linear regression on the data points are recorded on the left top.

Intrigued by this observation, we built a theoretical framework to better understand the origins of hashtag noise. We consider all cells positive for a given hashtag to be equivalent to a mathematical set. Since hashtag signals interfere with each other (Fig. 1C), these sets can potentially overlap with each other (Fig. 2C). As the number of such sets increase, the area of overlap is expected to increase, with a parallel decrease in the exclusive parts of each set, unless all higher order intersections (common overlap regions of multiple sets) tend to accumulate on the same cells. We derived a closed-form expression to calculate the exclusive regions (*E*) of a set among multiple interacting sets (Fig. 2D, Text S1) based on the simplifying assumption that the intersections are symmetric. This closed-form expression depends on three variables: pair-wise overlap *x*, higher-order interaction parameter *y* and the number of sets *n*. Note that *y* ≥ 1 and higher values of *y* means that the positive signals for multiple hashtags are increasingly not found in the same cells. Calculating *E* with arbitrary values of *x, y* and *n* (charts, Fig. 2D) with increasing *x, y* and *n*, confirms that the exclusive signal decreased rapidly.

Next, we estimated *x* and *y* of the hashtags in our pools (Fig. 2E, F, Methods) by randomly selecting hashtags and calculating *x* as the percentage of cells which are positive for the given hashtag and rest of the hashtags in the pool. The estimates of *y* was always lower in pools with *n* = 4 (Fig. 2E) when compared to the estimates of *y* in pool-F (Fig. 2F). Our model fitting confirms that with increasing number of hashtags, the fraction of singlets keeps decreasing because of the non-overlapping lower order intersections, i.e., rapidly diminishing higher order intersections.

Next, we fitted the %gap between GDE and 100 to linear regression (Fig. 2G) and observed a highly significant positive slope. The latter indicated that for every hashtag added to a pool, double the percentage of cells will be lost (e.g., 14% cell loss for 7 hashtags). Currently, wells can be super-loaded with up to 24 hashtags.

Hashtag contamination and capturing cells in droplets during emulsification are stochastic molecular processes. Hence, the overlaps of such interactions are expected to get compartmentalized, resulting in the loss of exclusive signals with an increase in the number of samples. Hence, higher estimates of *y* in pool-F are perhaps expected from a chemical kinetic perspective, too.

Due to cost considerations, we could not include compound design experiments with pools with intermediate (4 < *n* < 12) or very high (12 < *n* < 24) numbers of hashtags, though using an external dataset helped us partially address such gaps (Fig. 2G). The other limitation is that moderating designs are useless for the literature-based atlasing activities [17-19]. However, for labs planning experiments for building an atlas – moderated designs we discussed here can be useful to maximize the cell-yield and yet gracefully integrate the data. It has not escaped our notice that our ‘baseline’ chain design might facilitate post-integration, statistical finetuning for biological signal preservation across batches [20]. While our manuscript is based on a CITE-seq experiment, we speculate that our conclusions should be generally applicable for any other droplet based single-cell sequencing technique that involves sample multiplexing through hash-tagging [8, 10, 21].

## Conclusion

This rubric of moderated designs can be suited for diverse experimental requirements – from high-multiplexed clinical samples to low/non-multiplexed, referenced atlas building. The reference or the chain designs of multiplexing reduces the number of hashtags being used while mitigating batch-effects through integration of the pools. Moreover, in reference design, we can identify a threshold for the extent of biological mixing targeted in the samples, and then fine-tune integration strategies till such a level of mixing is achieved. A confounded multiplexing design, although more convenient to carry out for serially incoming samples, will be more challenging for batch-effect correction.

## Supporting information

Supplementary Material (Chatterjee et al)

## Acknowledgement

The work was supported by funding from NIAID (1R01AI168192) and DARPA (HR001121S0037-AIM-FP-009) to N.J. Singh.

## Methods

### Data generation for use in analysis

The experimental data used in this analysis is derived from an ongoing analysis of mouse splenocytes treated *in vivo* with five different variations of plasmid-encoded cytokines of the interleukin-12 family, compared to a control plasmid (Gorga, K. et. al, unpublished). The biological experiment involved the injection of expression constructs of the IL-12 protein family as plasmids in mice using hydrodynamic injection. Different versions of plasmids expressing IL-12P40 or variations therein were used – but not discussed here since the ensuing phenotypes are not studied here. Notably, we carefully controlled well-known factors contributing to batch-effects in our experiment: reagents, instruments, same-day execution, and same protocol for the sample processing, library preparation and sequencing. We skipped the mouse hashtags 10 and 11 (BioLegend) in our designs here since in one of our previous experiments [22] we observed that the reads of these two hashtags are highly correlated among each other across several pools (Fig. S6).

### Sequencing and data extraction

The pooled libraries were sequenced in the Illumina Novaseq platform (Psomagen Inc. Rockville, MD) and the base call (BCL) files demultiplexed with mkfastq function of CellRanger (10x genomics), to generate FASTQ files containing the reads of different pools.

### Alignment, data cleanup and processing

Both transcript and surface reads from each pool (i.e., index FASTQ files) were aligned with the count function from CellRanger, in default settings. The filtered barcode matrices (the output from CellRanger) were further analyzed with Seurat (v5.3.0) toolkit in R (v4.5.0). Six pools (A-F) were cleaned (count < 20,000, 500 < feature < 3,000, mitochondrial reads < 2.5%) and demultiplexed with the default Seurat pipeline (HTOdemux function). We removed the TCR, BCR, ribosomal and mitochondrial genes before integration since the variability within these sequences are a confounding factor independent of the design considerations of this manuscript. After cleanup we had a total of 44,846 cells from six pools (i.e., design-I).

### Integration of data designs

For a detailed description of the data designs see figure 1D. We processed the data in each design through three computational steps: transformation, integration across pools and measurement of batch-effects. For the analysis of scRNA-seq datasets, currently there are 22 different methods for transformation [23], 21 unique methods for integration [1, 4, 24] (can be applied with or without highly variable gene [HVG] selection and scaling as the preprocessing step) and 8 different ways of evaluating batch-effects [2]. Although there are several tool benchmarks available for these different steps [25], evaluation of the best pipeline, which is a combination of three steps would be computationally prohibitive (22 transformations × 4 preprocessing options × 21 integrations × 6 designs × 8 batch-effect calculations). Instead, we identified single cell benchmarks based on the literature and used consensus tools for these different steps (tools that performed well across different benchmarks, across diverse datasets and conditions). For transformation we chose shifted log transform and SCtransform [23, 26]. For integration methods, benchmarks are unclear [1, 4, 24], but we use harmony [27], CCA [28] and RPCA [29] ‘anchor’ integrations. Of course, there are other tools with similar potential performance capabilities that are not considered here in the interest of time and effort [14, 30, 31] (Fig. S1, S2A). We used a shifted log transformed (‘log-transformed’ in the text) but unintegrated version for all designs (as the comparators against the integrations) except design V since this involves just one pool (i.e., did not require an integration step) but was log transformed followed by PCA (see Methods).

For all designs, the following steps were taken for the transformation and integration: after merging the data from different pools in the given design, first the counts were normalized with Seurat function NormalizeData. This function carries out the log-transformation [23]. Subsequently, PCA was carried out after scaling and finding 3000 most variable features, followed by cell clustering and projection (UMAP). This is unintegrated version of the designs. Next, for all designs, except design-V, the following integration recipes were run on the latter principal component space: CCA [28], RPCA [29] and Harmony [27], utilizing the IntegrateLayer function from Seurat. Further cell clustering and projecting were carried out using each of these integrated spaces. PCA was always on first 50 dimensions and resolution parameter of cluster search (Louvain method) was always 1. A similar integration pipeline was carried out next but now on the SCtransformed data [26].

### Measurements of batch-effects

Conventionally, different pools/wells are called batches (i.e., the instances of separate library prep). However, the pooled samples may also have ‘batch’ effects apart from their expected biological variations due to possible differences in sample preparation steps. The integration methods are intended to minimize both types of batch-effects [3, 28, 32]. We, therefore, measured both types of batch-effects (i.e., pool and sample) in all designs and compared the entropy distributions of integrated and unintegrated outputs for the designs. These were then compared to the cell-cluster entropies, as a control group (Fig. 1B and 1E, F, bottom panels).

We used normalized Shannon entropy [2, 4] (‘entropy’ in the text for brevity) for the characterization of batch-effects. Briefly, Shannon entropy quantifies the information content in the neighborhood of each cell, following the expression below:

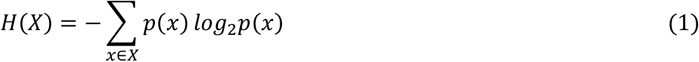

Where *X* is a random variable and *p*(*x*) is the probability of outcome *x* [33]. Let us assume, for example, that in design-I with six pools, in a neighborhood of a given cell (*k* cells) we have *a* cells from pool A, *c* from pool C, *e* cells from pool E (i.e., *a* + *c* + *e* = *k*). Hence the ‘pool entropy’ of that specific cell will be:

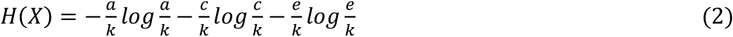

Hence entropy in this context is the average uncertainty of finding cells of a specific pool in the neighborhood. We used CellMixS package [2] in R for the calculation of entropy, where a normalized Shannon entropy (*H*_n_(*X*)) is calculated as follows:

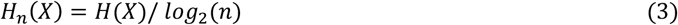

Where *n* is the number of categories within *X*. We used the evalIntegration function in CellMIxS package to this end. The normalized Shannon entropy is a unitless quantity that ranges between 0-1. The unintegrated or integrated spaces (for the different designs, integration recipes, transformations) were transferred as is to the latter function for the calculation of normalized Shannon entropies (through conversion of the Seurat objects to SingleCellExperiment objects [34] before using them with the evalIntegration function). Since the total number of pools, samples or cell-types (i.e., the categories, *n*) are very different in diverse designs, the neighborhood size, i.e., the *k*-nearest neighbors (*k*), should be adjusted for a fair comparison between the entropies measured for different metadata. We have set the neighborhood size (*k*) at three times the number of unique categories (*n*) for all our entropy calculations. For example, we have six pools in design-I, so we set *k* at 18 for pool entropy calculations in design-I, but to measure the cell-type entropy in design-I, in Harmony integration space, we set *k* at 78 (*n* for cell clusters in Harmony space = 26). Note that *H*_*n*_(*X*) is indeterminate when *n* = 1.

### Differential expression analysis across wells

Differential expression analysis was carried out (Fig. S3) using DESeq2 package [35] in R, after pseudo-bulking the counts at the hashtag level. The differentially expressed genes were checked for enrichment of biological functions using ShinyGO [36].

### Calculations of hashtag efficacy

Hashtag demultiplexing was carried out using the HTOdemux function in Seurat with parameters set to default [12, 37, 38]. Thresholds declared by HTOdemux were used further to calculate the GDE and IHE (Fig. 2A, B). The latter thresholds were also utilized for enumerating the pair-wise overlap parameter, i.e., *x* (Fig 2C-F, Supplementary text S1). Our closed form expression cannot be explicitly solved for *y* when *n* > 3. Since *n* ≥ 4 in our designs, we numerically estimated *y* from known *x, n* and *E. y* was estimated using the uniroot function in R for root search within predefined boundaries. For several values of *x*, the estimated values of *y* are noted in the figure, wherever it could be estimated within the identical fixed intervals for any hashtag in any pool.

### Using external data

We used one external dataset [7] for gathering more GDE data points in our regression analysis (Fig. 2G). This dataset (GSE108313) was made available in the convenient .rds format by the authors as part of the vignette for the Seurat HTOdemux algorithm. The correlogram of the hashtag reads (Fig. S6) was prepared using one of our own datasets [22] published previously (PRJNA1188714).

## Data Availability

All code for generating all main text and supplementary figures are available as a GitHub repository: https://github.com/NevilLab/Hashbatch

## Notes

### Competing Interest Statement

The authors have declared no competing interest.

