## Supplementary Material (Chatterjee et al) for "Moderated designs can balance between batch-effect mitigation and cell loss due to hashtag-assisted pooling in single-cell experiments"

**Title:**

Running title: Designs for balancing batch-effect and cell loss

**Authors:**

Budha Chatterjee[1], Katrina Gorga[1], Carly Blair[1], Yuko Ohta[1], Elizabeth M. Hill[1], Christopher T. Boughter [2], Martin Meier-Schellersheim [2] & Nevil J. Singh[1 *]

This file contains the following items:

Figure S1 (page 2-3): Log transformed (LN) or SCtransformed (SCT) datasets for different designs and integrations showing the pools.

Figure S2 (page 4-5): Projected datasets depicted to show sample mixing.

Figure S3: DEGs in the same sample across wells or different samples shown with volcano plots.

Figure S4: The distribution of the entropies in the designs across the integration recipes and transformations.

Figure S5: Distributions of counts and features in doublets (Do), negatives (Ne) and singlets (Si).

Figure S6: A correlogram of hashtags 1 to 14 (gray bars at the top and left) for their CLR normalized reads.

Text S1: Derivation of a closed form expression to represent exclusive parts of sets (i.e., hashtag positive events) with higher order interactions.

25

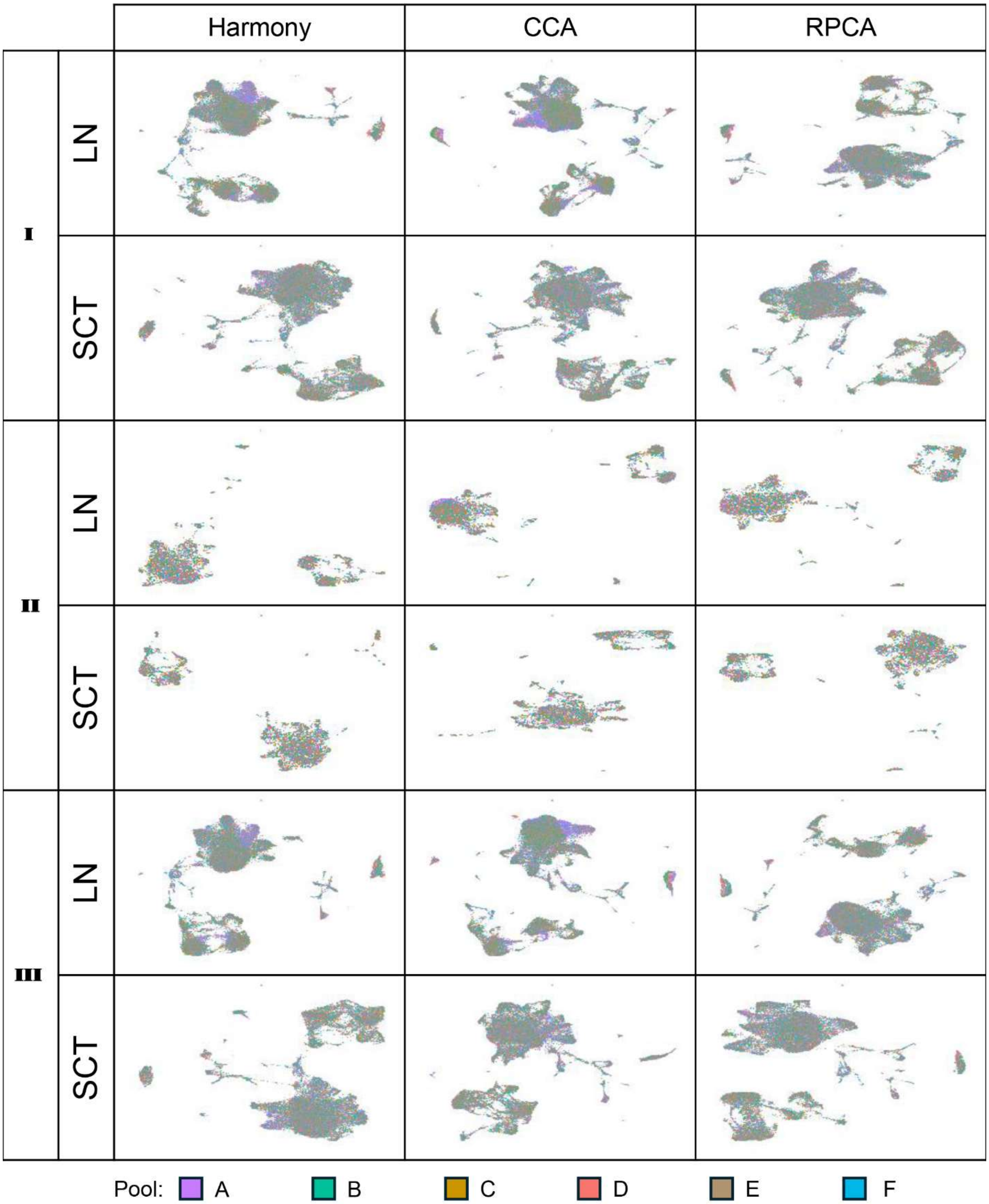

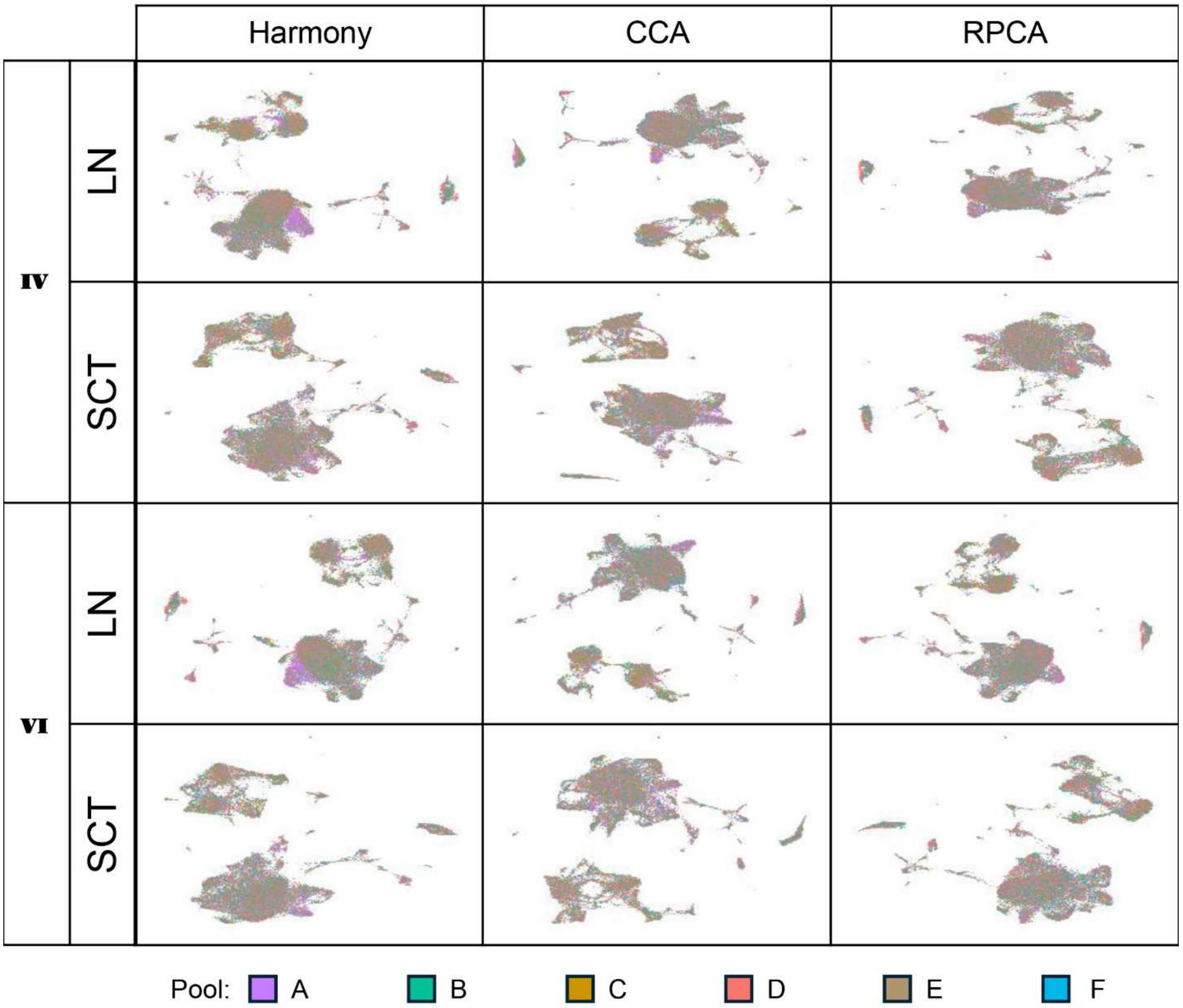


**Figure S1 (page 2-3): Log transformed (LN) or SCtransformed (SCT) datasets for different designs and integrations showing the pools.** UMAPs of all combinations of designs, transformations and integration recipes are shown for the pools. The colors of the cells from specific pools are provided at the bottom. Design V is not shown since it contains only one pool (pool F) and hence not integrated.

32

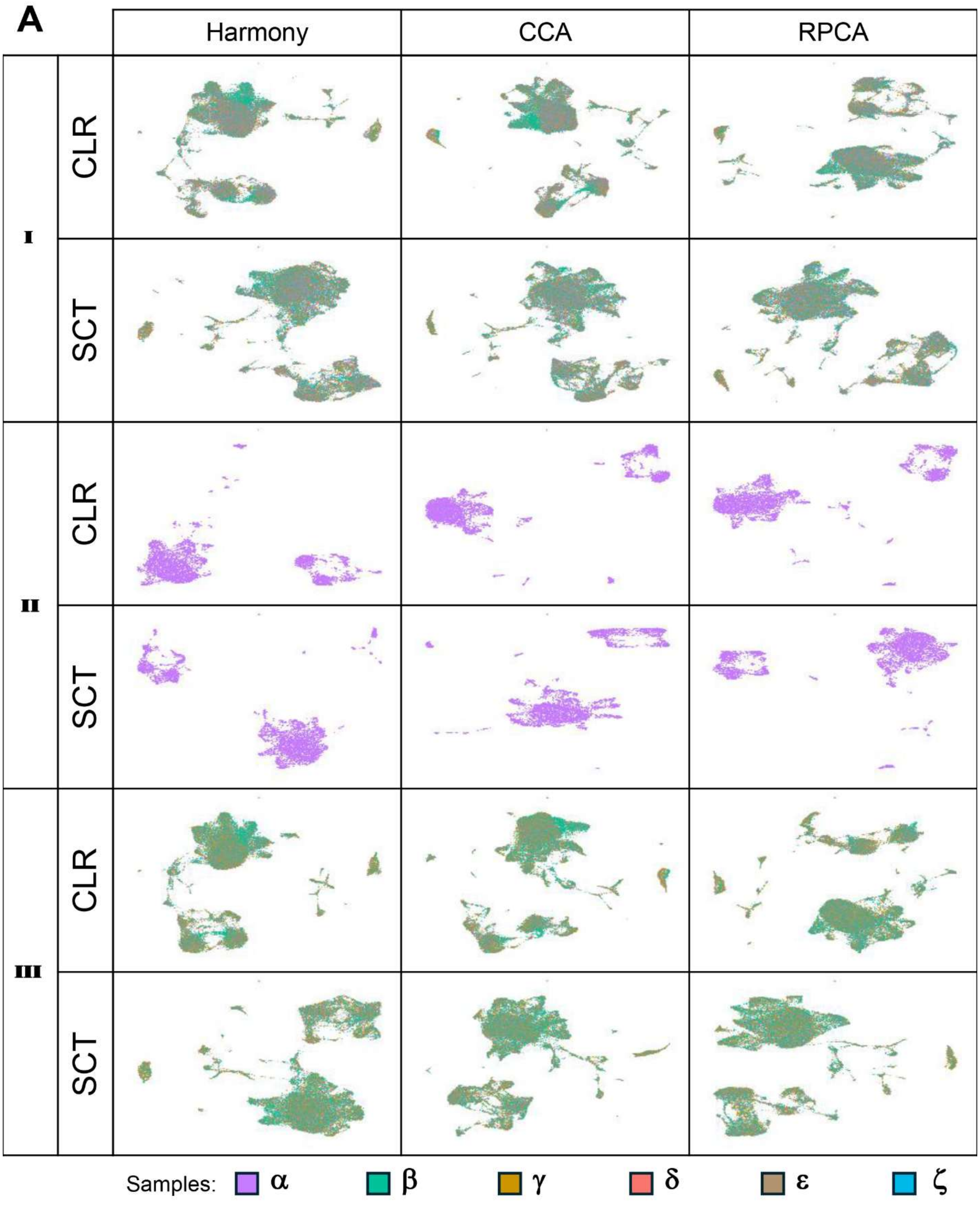
A
Harmony
CCA
RPCA
I
CLR
SCT
II
CLR
SCT
III
CLR
SCT
Samples: α β γ δ ε ζ

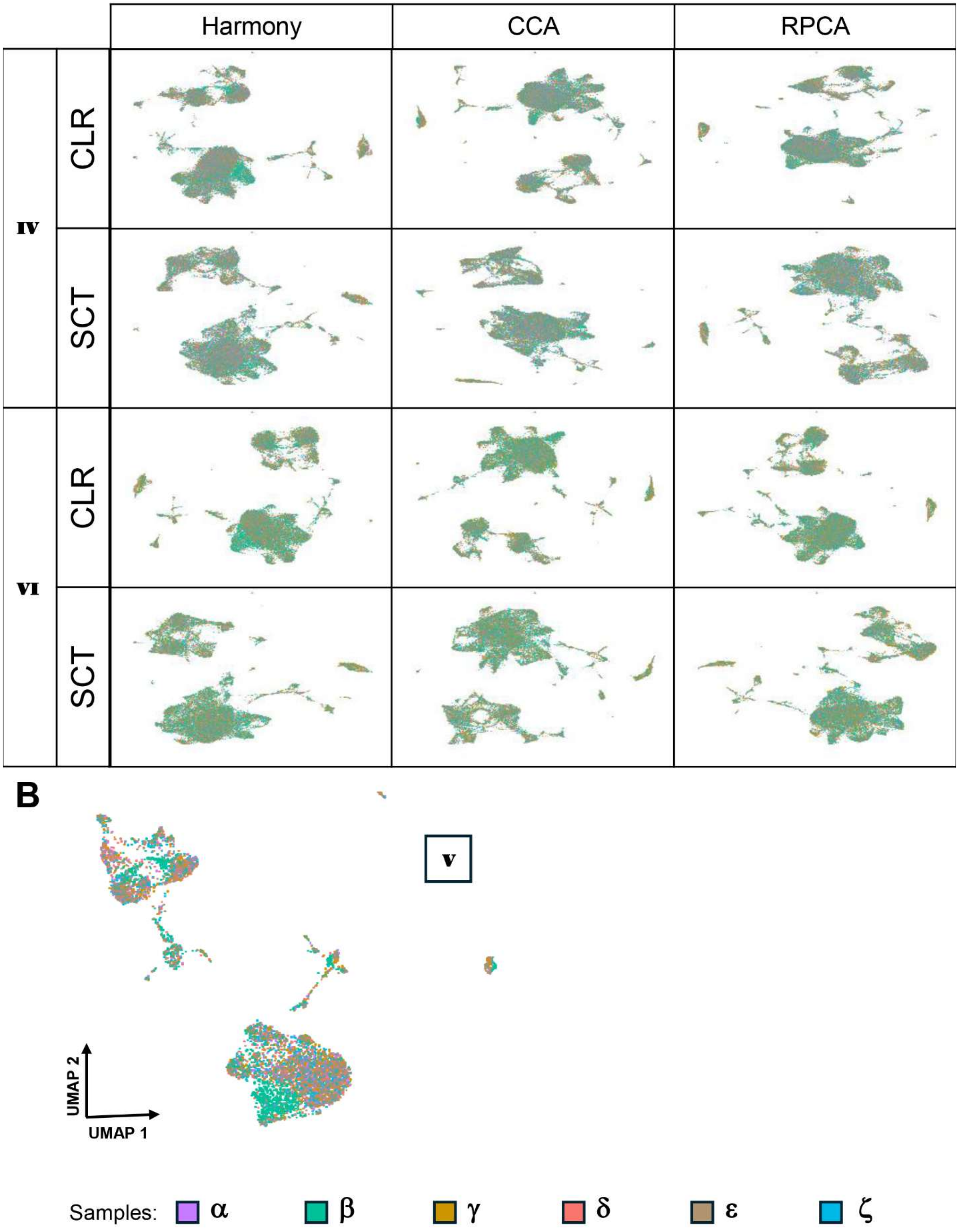

Harmony
CCA
RPCA
iv
CLR
SCT
vi
CLR
SCT
B
v
UMAP 2
UMAP 1
Samples: α β γ δ ε ζ

**Figure S2 (page 4-5): Projected datasets depicted to show sample mixing.** (A) Same as previous supplementary item but now showing the mixing of the samples. The colors of the cells from different samples are shown in the bottom. (B) The unintegrated log transformed design V, the UMAP on the PCA space is shown to depict within well mixing of the samples.

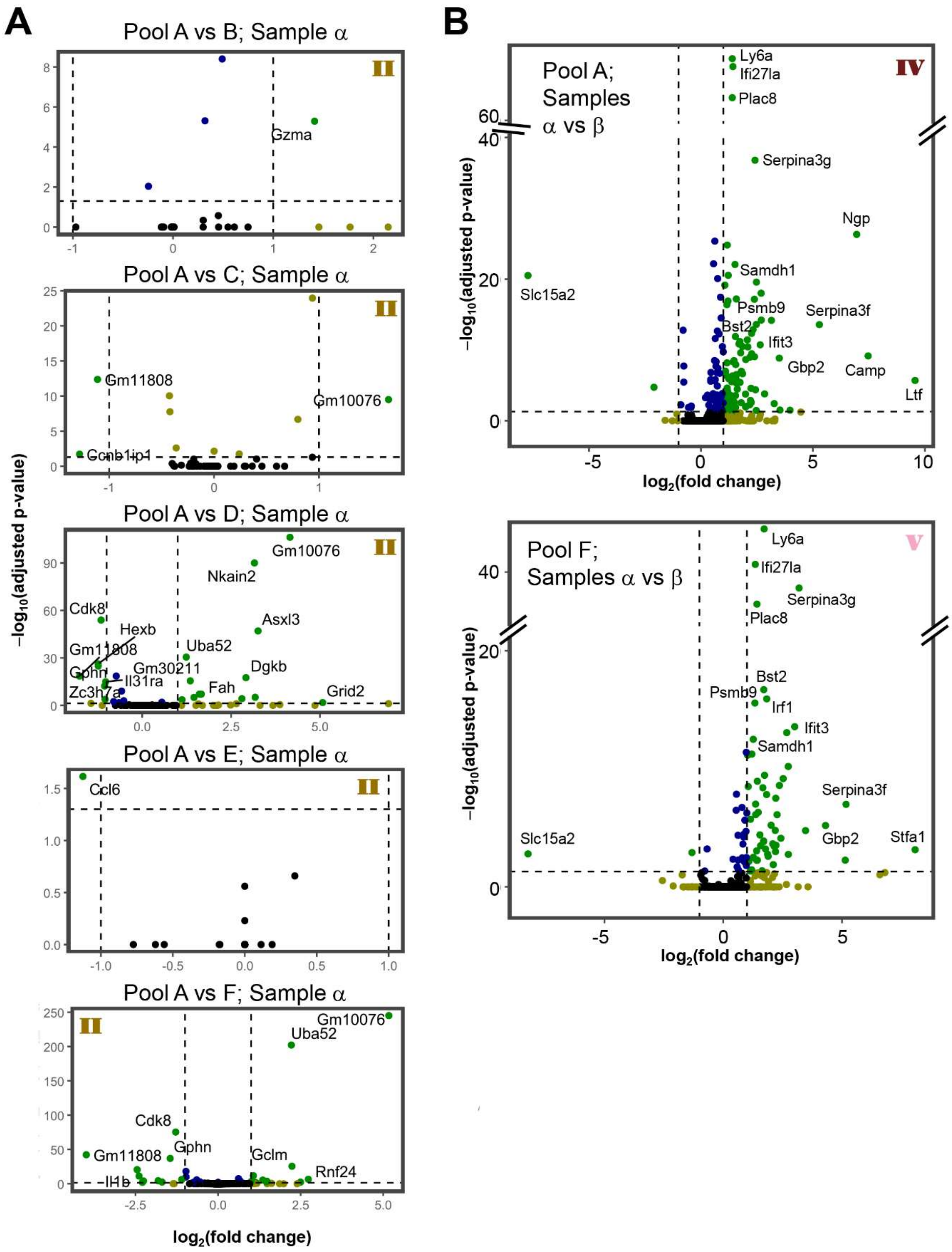


**Figure S3: DEGs in the same sample across wells or different samples shown with volcano plots:** (A) comparison of sample α in well A vs rest of the wells (design II). AvsB, AvsC and AvsE had one or two disparate genes up/down regulated. AvsD and AvsF showed upregulation of 13 and 11 genes respectively among which 3 were common. These genes (13, 11 in AvsD, AvsF respectively) did not show any function enrichment at a significant level (p= 0.01). (B) Comparison of different samples such as α vs β from different pools – pool A and pool F from designs IV and V respectively. 55 and 95 genes are upregulated in pool A and F respectively – 47 of them are common in both. Visual inspection of the volcano plots indicate shows the common genes with largest statistical or numeric significance.

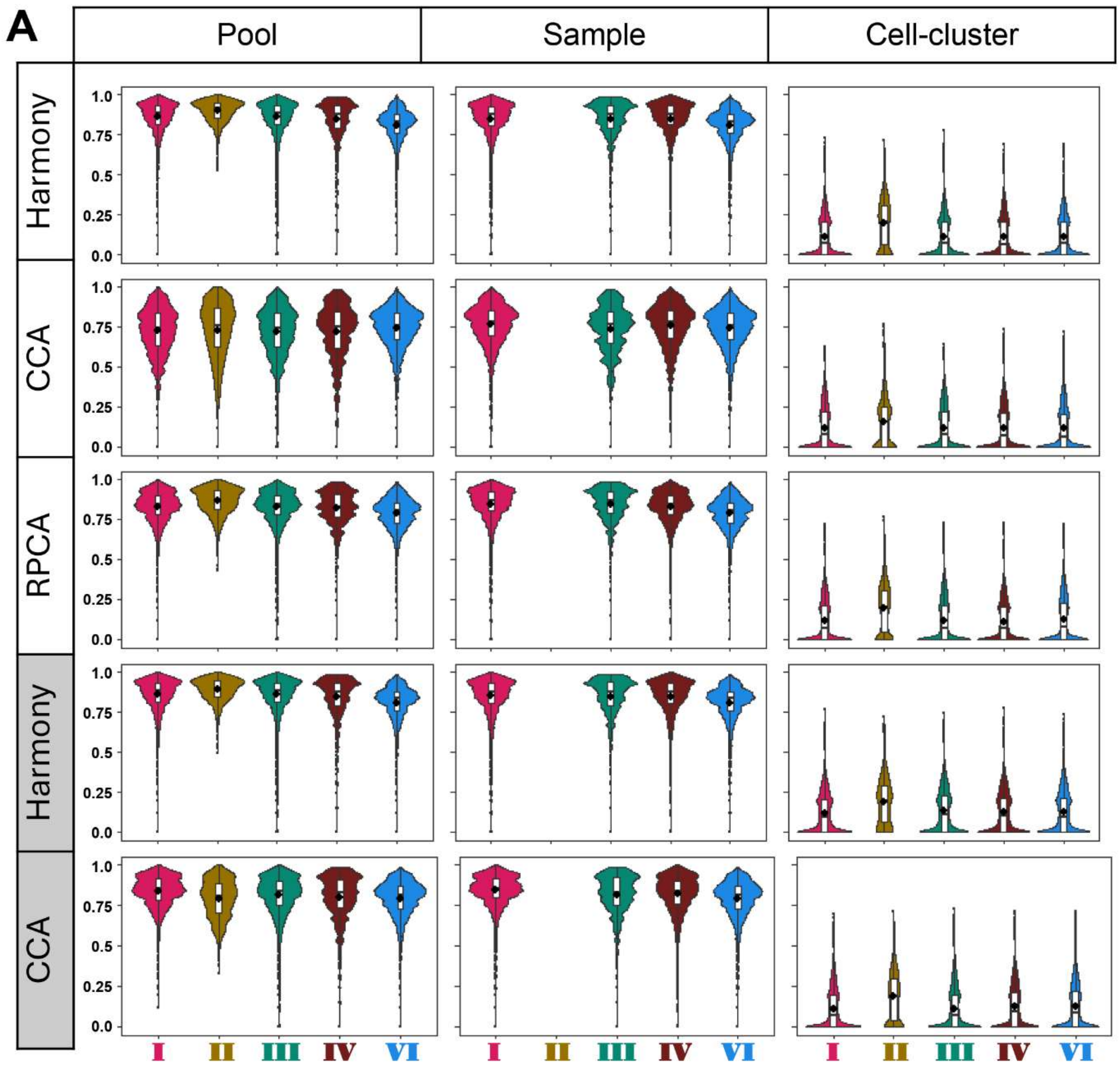


**Figure S4: The distribution of the entropies in the designs across the integration recipes and transformations.** The gray boxes represent SCtransformed data. Pool, sample and cell type entropies are shown as in the maintext figures 1E and F.

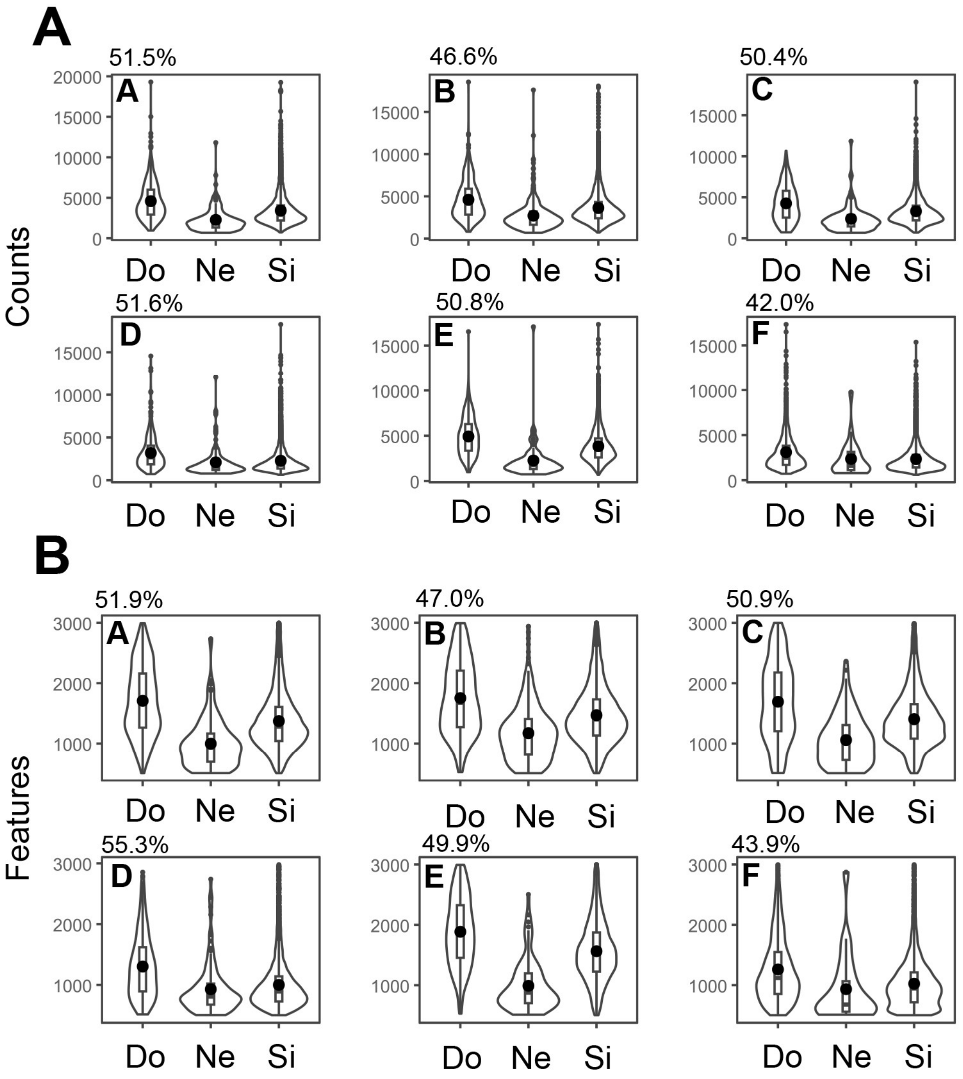


**Figure S5: Distributions of counts and features in doublets (Do), negatives (Ne) and singlets (Si):** (A) Distributions of the counts in doublets, singlets and negatives in six different wells: from A to F. The well id is indicated at the top left inside each plot. The percentage at the top is the percentage of cells in doublets below the $3^{rd}$ quartile of the singlets. (B) Same as (A) but distributions represent features (i.e., number unique genes in each cell) now.

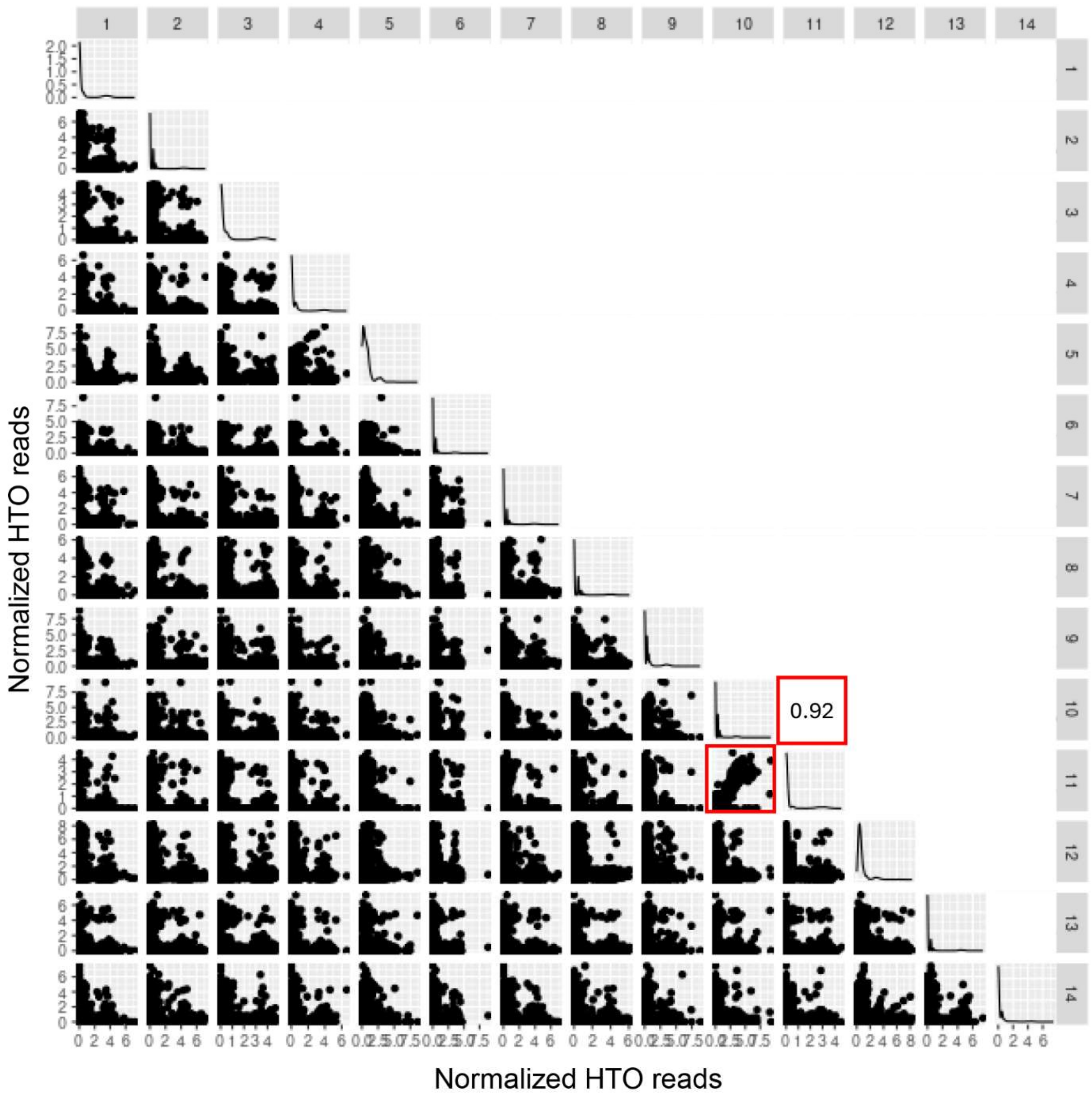


**Figure S6: A correlogram of hashtags 1 to 14 (gray bars at the top and left) for their CLR normalized reads.** Only the normalized read counts hashtag 10 and 11 showed a correlation coefficient above 0.1. All other (numbers not shown) combinations showed correlation below 0.1.

**Text S1: Derivation of a closed form expression to represent exclusive parts of sets (i.e., hashtag positive events) with higher order interactions.**

Let's assume two sets have an intersection $x$, and this pair-wise intersection remains constant for all binary combinations of $n$ sets. Next, let's assume that $y$ is the higher order intersection parameter, such that, say for $n = 3$, the extent of ternary intersection is $x/y$, and for $n = 4$, the quaternary intersection is $x/y^2$, and so on. So, for $k + 1$ sets we can generalize intersection ($I_{k+1}$) as follows:

$$I_{k+1} = \frac{x}{y^{(k-1)}} \qquad where\ k \geq 1 \tag{1}$$

Now, let's further assume that we are to calculate the exclusive region ($E$) of a set $A$, which intersects with $n$ different sets, and these sets participate in higher order interactions symmetrically. To that end, we can use the Inclusion-Exclusion Principle (IEP) [1, 2], having the total area (i.e., a representation of all cells positive for a given hashtag) of $A$ as $|A|$:

$$E = |A| - \binom{n-1}{1} I_2 + \binom{n-1}{2} I_3 - \binom{n-1}{3} I_4 + \cdots + (-1)^{n-1} \binom{n-1}{n-1} I_n \tag{2}$$

Now following (1), and assuming that total area of set $A$, i.e., $|A|$ is 1, we can replace the interaction terms ($I_{k+1}$) as follows:

$$E = 1 - \binom{n-1}{1} x + \binom{n-1}{2} \frac{x}{y} - \binom{n-1}{3} \frac{x}{y^2} + \cdots + (-1)^{n-1} \binom{n-1}{n-1} \frac{x}{y^{n-2}} \tag{3}$$

A convenient, specialized identity of the IEP is as follows:

$$E = |A| - \sum_{k=1}^{n-1} (-1)^{k-1} \binom{n-1}{k} I_{k+1} \tag{4}$$

Replacing $I_{k+1}$ following (1) and (3),

$$E = 1 - x \sum_{k=1}^{n-1} (-1)^{k-1} \binom{n-1}{k} \left(\frac{1}{y}\right)^{k-1} \tag{5}$$

Now, following the binomial theorem, the expansion of $(1 - r)^m$ is as follows:

$$(1 - r)^m = \sum_{k=0}^{m} \binom{m}{k} (-r)^k \tag{6}$$

Following simplification and rearrangement:

$$\frac{1 - (1 - r)^m}{r} = \sum_{k=1}^{m} (-1)^{(k-1)} \binom{m}{k} r^{(k-1)} \tag{7}$$

Replacing $r = 1/y$ and $m = n - 1$ and substituting (5):

$$E = 1 - xy + xy\left(1 - \frac{1}{y}\right)^{(n-1)} \tag{8}$$

Further simplification yields:

$$E = 1 - xy + x\frac{(y-1)^{(n-1)}}{y^{(n-2)}} \tag{9}$$
